# *In situ* architecture of the algal nuclear pore complex

**DOI:** 10.1101/232017

**Authors:** Shyamal Mosalaganti, Jan Kosinski, Sahradha Albert, Miroslava Schaffer, Jürgen M. Plitzko, Wolfgang Baumeister, Benjamin D. Engel, Martin Beck

**Author notes:** contributed equally (Shyamal Mosalaganti, Jan Kosinski, Sahradha Albert, Miroslava Schaffer).

## Abstract

Nuclear pore complexes (NPCs) span the nuclear envelope and mediate nucleocytoplasmic exchange. They are a hallmark of eukaryotes and are deeply rooted in the evolutionary origin of cellular compartmentalization. NPCs have an elaborate architecture that has been well studied in vertebrates. Whether this architecture is unique or varies significantly in other eukaryotic kingdoms remains unknown, predominantly due to missing *in situ* structural data. Here, we report the architecture of the algal NPC from the early branching eukaryote *Chlamydomonas reinhardtii* and compare it to the human NPC. We find that the inner ring of the *Chlamydomonas* NPC has an unexpectedly large diameter, and the outer rings exhibit an asymmetric oligomeric state that is unprecedented compared to all previously proposed models of NPC architecture. Our study provides evidence that the NPC is subject to substantial structural variation between species. The divergent and conserved features of NPC architecture provide insights into the evolution of the nucleocytoplasmic transport machinery.

## Introduction

Nuclear pore complexes (NPCs) mediate molecular traffic between the cytoplasm and nucleus, and are therefore indispensible for eukaryotic life. NPCs are built from ~30 nucleoporins (Nups) that are mostly conserved across eukaryotes, with some exceptions^1-3^. Nups are organized into various subcomplexes, which assemble together to form two outer rings that reside in the cytoplasm and nucleus, and an inner ring that fuses the inner and outer nuclear membranes. In the human NPC (*Hs*NPC), the ten-membered Y-complex is a major component of the outer rings (also referred to as the cytoplasmic and nuclear rings). 32 copies of the Y-complex arrange in a head-to-tail conformation to form concentric, reticulated rings within both the cytoplasmic and nuclear rings^4^. The Y-complex scaffold is complemented by additional subcomplexes that fulfill specific functions in the nuclear and cytoplasmic periphery and provide the directionality cue for nucleocytoplasmic exchange. The inner ring is composed of 32 protomers, each containing the Nup93 and Nup62 subcomplexes. Although the inner ring is constructed from proteins different than the outer rings, the oligomeric assembly of the inner and outer rings is similar^5,6^.

This architectural model of the human NPC is based on *in situ* cryo-electron tomography (cryo-ET) and subtomogram averaging of NPCs imaged within isolated HeLa cell nuclear envelopes. Similar structural analysis is also available for intact HeLa cells^7^, u2os cells^8^ and *Xenopus laevis* nuclear envelopes^9^. Analyses of the *Dictyostelium discoideum^10^* and *Saccharomyces cerevisiae^11^* NPCs lacked the necessary resolution to visualize subcomplex architecture. Various biochemical and structural studies of NPC subcomplexes from vertebrates, fungi and Trypanosomes have concluded that the subcomplexes are conserved (for a comprehensive review see^12^). However, it remains unclear whether subcomplexes from different species assemble into NPCs in an identical fashion. This is highlighted by a prominent model proposed for yeast NPC architecture that suggests that yeast have fewer Y-complexes than humans^3^. Thus, the number of Y-complexes and oligomeric state of the NPC across eukaryotic kingdoms remains uncertain.

An important architectural feature underlying all previously proposed models of NPC architecture is the intrinsic C2 symmetry of the inner ring and Y-complexes across the plane of the nuclear envelope^1^·^3,12^. It has been proposed that the NPC’s remarkable degree of symmetry might be essential to facilitate the modular assembly of its large macromolecular structure from a limited set of building blocks^13^. Here, we combine focused ion beam thinning of vitreous frozen cells^14-16^ with *in situ* cryo-ET to analyze NPC architecture within the native cellular environment of *Chlamydomonas reinhardtii*, a unicellular green alga (Chlorophyte) and an early branching eukaryote. This approach facilitates structural analysis within intact cells in a close-to-live state without the need for subcellular fractionation or affinity purification. We find that the *C. reinhardtii* NPC (*Cr*NPC) has several distinct architectural features, including an asymmetrical oligomeric state of the cytoplasmic and nuclear rings. We conclude that different mechanisms of Y-complex oligomerization have evolved independently for the *C. reinhardtii* cytoplasmic and nuclear rings, and that NPC architecture may vary considerably throughout eukaryotic life.

## Results

### Key scaffolding subcomplexes are conserved in C. reinhardtii

*C. reinhardtii* cells are particularly well suited for *in situ* structural biology, enabling high-resolution imaging of cellular structures^17-21^. This model organism is therefore an excellent candidate to address the question of how well current models of NPC architecture are transferable across eukaryotic species. We first explored the genome of *C. reinhardtii^22^* by sequence alignments to determine whether the key Nups of the NPC are detectable in the genome and whether the Nup subcomplexes are conserved. In agreement with a previous genomics study^23^, we found homologs of all major scaffold and FG-Nups (Supplementary Fig. 1, Table 1). *NUP188* gene, which was previously reported to be absent in plants^24,25^, was present in the *C. reinhardtii* genome. We also detected a *NUP188* homolog in the genome of *Arabidopsis thaliana*, emphasizing that Nup188 protein has a conserved role in the NPC scaffold architecture and is likely an ancient protein. Although sequence similarity cannot prove that an individual gene indeed encodes a functional equivalent of another gene, it is fair to conclude that the inner ring and Y-complexes are generally conserved in *C. reinhardtii* because all components that have been functionally analyzed in various species^12^ were confidently detected.

However, we did not detect *NUP358* and *NUP153* genes, which in metazoa constitute cytoplasmic ring and nuclear ring-specific elements, respectively. The Y-complex member, *NUP37*, and the transmembrane Nups, *GP210* and *POM121*, are also absent from the genome, whereas the chromatin-binding Nup, Elys, is encoded in a truncated form. Failure to detect these genes might be due to low sequence similarity or insufficient sequencing coverage of the genome. However, in the case of Nup358, it has been well established that this protein has evolved in animals and is absent from fungi and plants^26^.

### The algal NPC has an unprecedented architecture

To analyze the *in situ* NPC architecture of *C. reinhardtii*, we acquired tomograms of the nuclear envelope within its native cellular environment (Supplementary Fig. 2b) and extracted 78 subtomograms containing individual *Cr*NPCs. We used subtomogram averaging to produce structural maps of the cytoplasmic, inner and nuclear rings at an overall resolution of ~3 nm (Supplementary Fig. 2a,c)^17^.

Comparison of the *Cr*NPC to the *Hs*NPC revealed striking differences in their overall dimensions and architecture (Fig. 1a). In humans, the outer rings are oriented in an upright position and are spatially separated from the inner ring by a connector element (magenta arrowheads, Fig. 1a)^27^. In *C. reinhardtii*, however, the outer rings are flatter and are directly stacked onto the inner ring. This direct engagement of inner and outer rings enforces a compact conformation of the *Cr*NPC; the *Cr*NPC scaffold extends only ~60 nm along the nucleocytoplasmic axis, whereas the human NPC spans ~80 nm. While the outer diameters of the *Hs*NPC and the *Cr*NPC along the plane of the nuclear envelope are similar, the inner diameter of the *Cr*NPC central channel is approximately 21 nm wider than that of the *Hs*NPC (Fig. 1b), suggestive of a modified inner ring arrangement. Lastly, the *Cr*NPC’s cytoplasmic ring has considerably less density than the nuclear ring. Such extensive asymmetric density across the nuclear envelope plane is surprising and has not been previously reported for NPCs in any other organism (Fig. 1c). Although the cytoplasmic ring contains less density overall, it has distinct features within densities protruding towards the central channel (black arrowheads, Fig. 1c).

**Figure 1.**
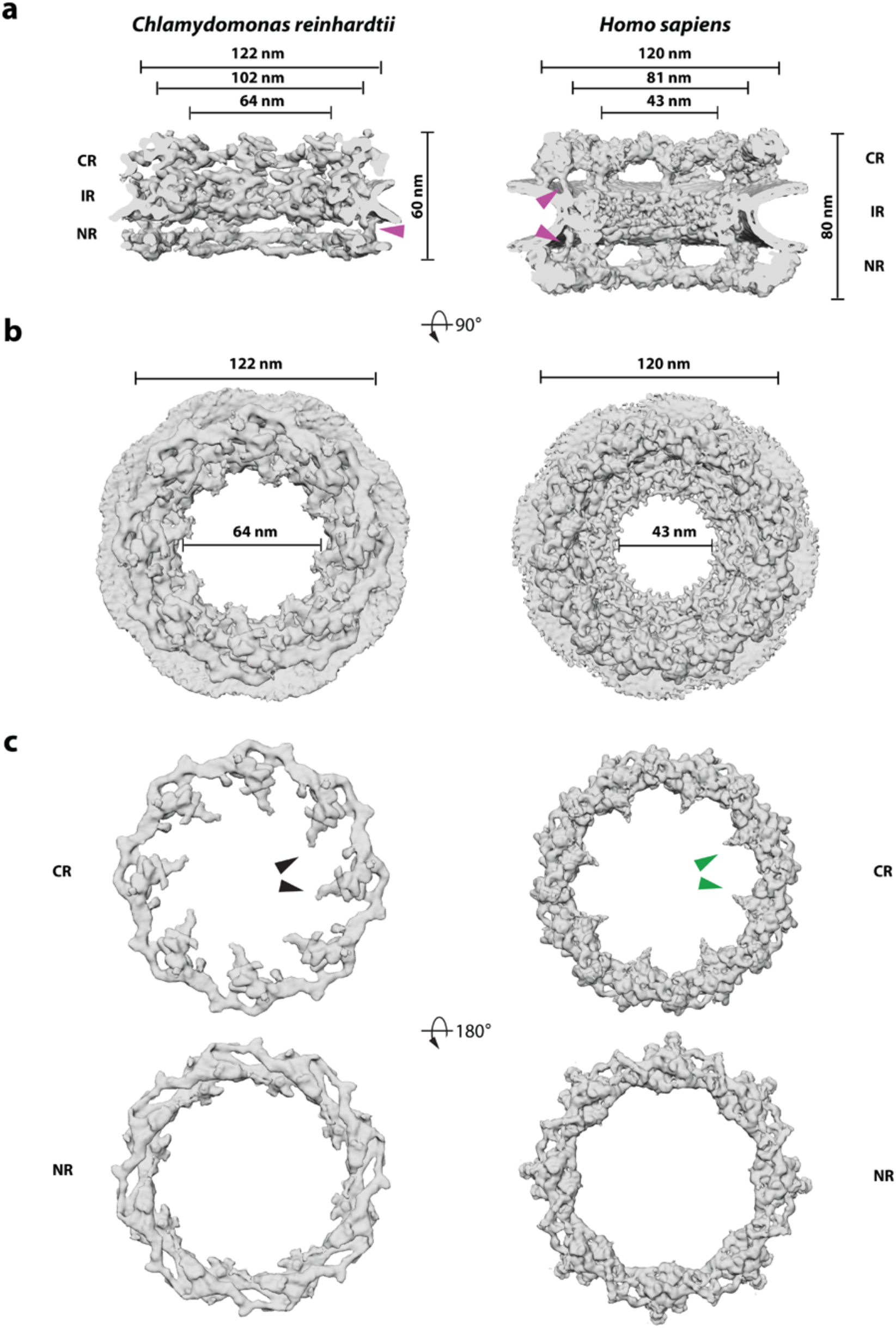
Structure of the *Cr*NPC in comparison to the *Hs*NPC. **(a)** Structures are displayed as rendered isosurfaces, sliced through the central axis. Magenta arrowheads indicate the connector element, which is absent from the cytoplasmic side of the *Cr*NPC. **(b)** Cytoplasmic face view. The dilation of the *Cr*NPC central channel is apparent. **(c)** Cytoplasmic and nuclear rings of the *Cr*NPC and *Hs*NPC. Black and green arrowheads indicate the density assigned to the Nup159 (Nup214 in humans) subcomplex, which forms cytoplasmic filaments that protrude towards the central channel. Abbreviations: cytoplasmic ring (CR), inner ring (IR), nuclear ring (NR).

### The algal inner ring is dilated but its basic organizational principles are conserved

We next assessed whether the architectural arrangement of scaffolding Nup subcomplexes that we previously assigned into the *Hs*NPC^4,5^ can explain the density observed for the subtomogram average of the *Cr*NPC. To this end, we used a hierarchical procedure that included an unbiased fitting of low-pass filtered structural models of human Y-complexes and inner ring protomers, filtering the fits using objective criteria (not clashing with each other and having at least 60% overlap with the map of the *Cr*NPC), and local re-adjustment of the selected fits to account for conformational differences between the *Cr*NPC and *Hs*NPC (Materials and Methods and Supplementary Fig. 3). The resulting density assignment reveals that the *Cr*NPC map can be well explained by the structural repertoire of human scaffolding Nups (Supplementary Figs. 4 and 5).

The 32 C2-symmetric protomers assigned into the *Hs*NPC inner ring^5,6^ not only fit the inner ring of the *Cr*NPC but also have an identical relative arrangement to that in the *Hs*NPC (Fig. 2a, Supplementary Fig. 4). The entire asymmetric unit, consisting of four C2-symmetric protomers, fits into the *Cr*NPC with high statistical significance (Supplementary Fig. 4a-c). Three out of four inner ring protomers were statistically significant after correction for multiple comparisons as assessed by systematic fitting; the one remaining protomer was recovered by subsequent filtering (Supplementary Fig. 4d-f). The identified density is weaker in the regions of the two inner protomers corresponding to the Nup62 subcomplex (Supplementary Fig. 4), leaving the exact number of Nup62 per asymmetric unit uncertain.

**Figure 2.**
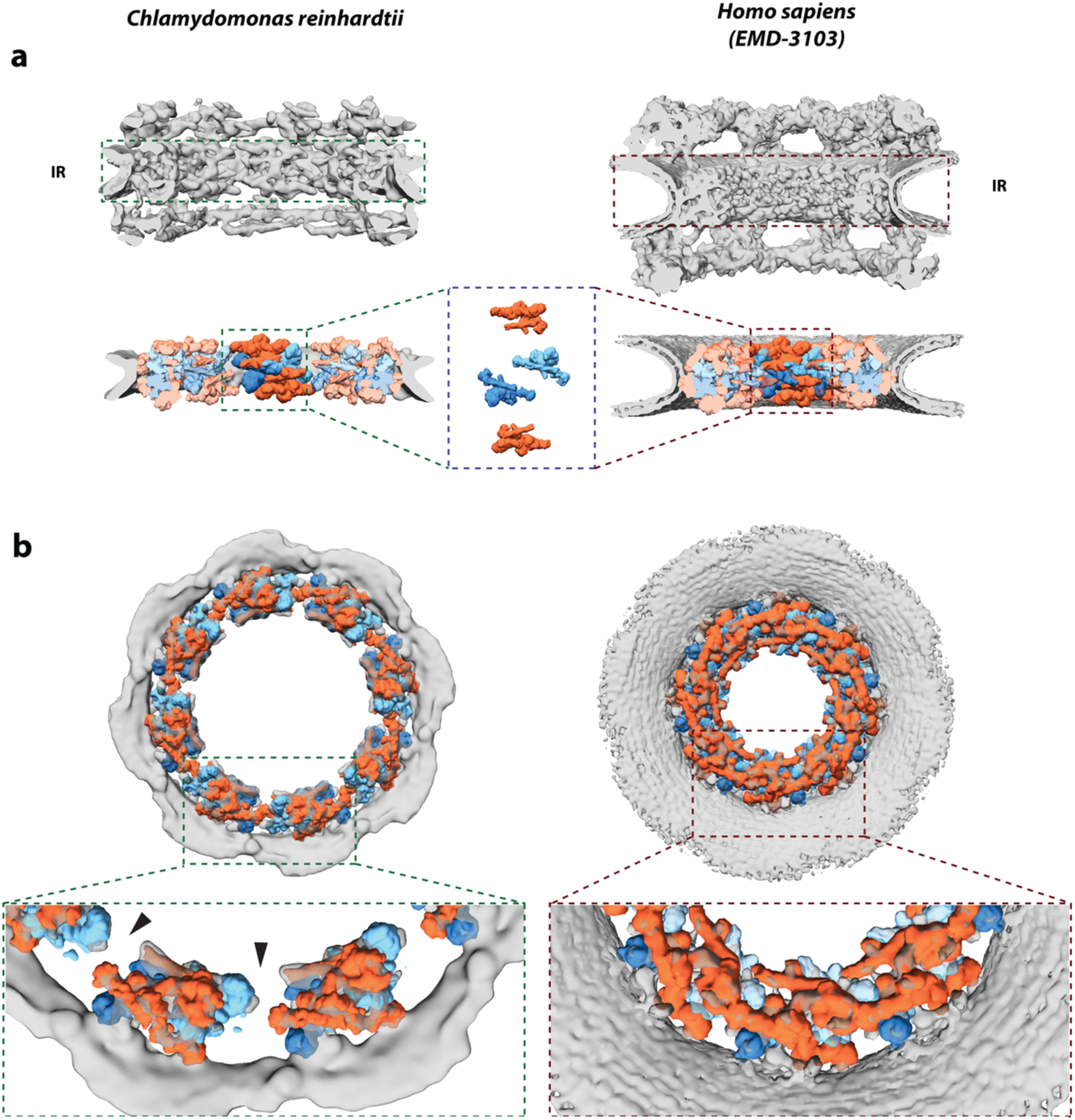
The inner ring of the *Cr*NPC is dilated compared to the *Hs*NPC. **(a)**Structures of the *Cr*NPC and the *Hs*NPC, displayed as rendered isosurfaces, sliced through the central axis. The inner rings are indicated with dashed boxes (top). The four protomers of the asymmetric unit (orange: outer protomers, blue: inner protomers), each containing Nup93 and Nup62 subcomplexes, explain the inner ring densities of both the *Cr*NPC (bottom left) and *Hs*NPC (bottom right). **(b)** View of the *Cr*NPC and *Hs*NPC inner rings seen along the nucleocytoplasmic axis. It is evident that the asymmetric units (spokes) of the *Cr*NPC inner ring are separated from each other (left), leaving rather large peripheral channels (arrowheads), whereas the asymmetric units of the *Hs*NPC are positioned closer together (right).

The four stacked protomers within the asymmetric units of the inner ring (traditionally termed spokes) are arranged with respect to each other in a similar fashion to the *Hs*NPC (Fig. 2a). However, the eight spokes of the *Cr*NPC are positioned in a wider arrangement, leading to an apparent dilation and wider central channel diameter (Fig. 2b). The tight interconnection between spokes observed in the *Hs*NPC is therefore relaxed in the *Cr*NPC, leading to gaps between the spokes that correspond to larger peripheral channels (black arrowheads, Fig. 2b). We conclude that although the principle composition and architecture of the inner ring within each asymmetric unit is conserved between these two distantly related eukaryotes, the overall spacing of the spokes is strikingly different.

### The cytoplasmic ring of the algal NPC has a simplified oligomeric state compared to the human NPC

We next examined the outer rings in detail. In humans, it has been established that the Y-complexes account for the majority of the observed outer ring density. In both the cytoplasmic and nuclear rings of the *Hs*NPC, 16 Y-complexes oligomerize in head-to-tail fashion to form reticulated double concentric rings^4^ (Fig. 3a). We identified Y-complexes at the expected positions in the *Cr*NPC map (Fig. 3a) using exhaustive fitting of low-pass-filtered human models, albeit with lower scores compared to the inner ring fitting, i.e. the assignments do not rise to statistical significance during the exhaustive fitting but are identified by the subsequent filtering step (Supplementary Figs. 3 and 5). This may be attributed to the hinges within the Y-complexes^28^, which appear to adopt a different conformation in the *Cr*NPC. However, the characteristic Y-complex shape is obvious in the map of the *Cr*NPC (Fig. 3a, Supplementary Fig. 5, Video 1). In both the cytoplasmic and nuclear rings of the *Cr*NPC, the Y-complexes are tilted down towards the dilated inner ring, resulting in flatter outer ring architecture than in the *Hs*NPC.

**Figure 3.**
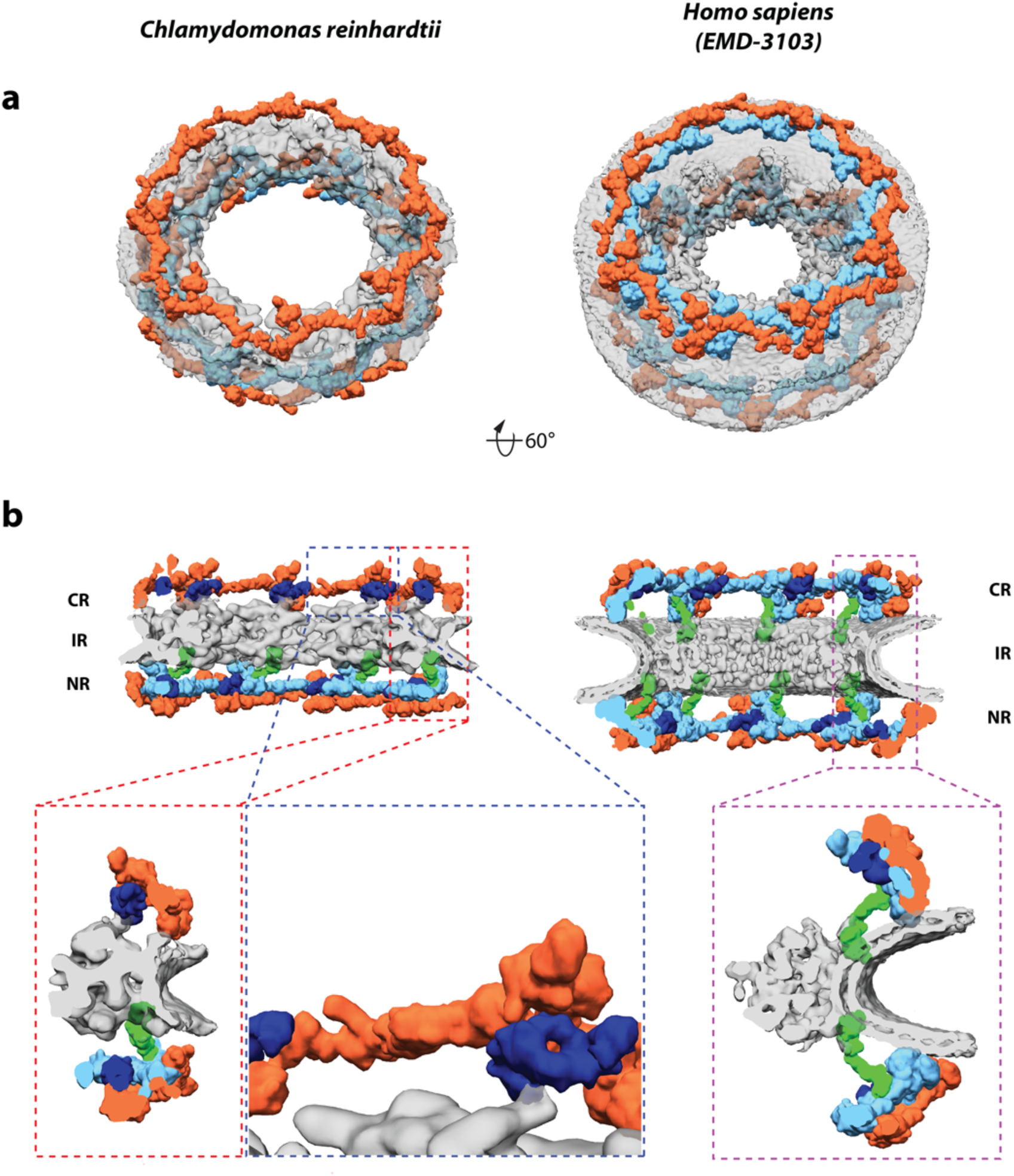
The *Cr*NPC has 24 Y-complexes. **(a)** Segmented Y-complexes according to the fits presented in Supplementary Figs. 3 and 5 are shown superimposed with the inner ring structure (grey). The distribution of Y-complexes in the *Cr*NPC is asymmetric across the nuclear envelope plane. The cytoplasmic ring has only 8 Y-complexes (orange), whereas the nuclear ring has 16 (orange and light blue). In the *Hs*NPC, the distribution is symmetric, with 16 Y-complexes in both of the outer rings. **(b)** Rotated views of the *Cr*NPC and *Hs*NPC, sliced through the central axis and colored as in panel **a**. Density attributed to large scaffold Nups (Nup205/Nup188) in the outer rings of the *Hs*NPC) (dark blue between the inner (bright blue) and outer (orange) Y-complex. Similar density is observed in the *Cr*NPC, although this assignment remains tentative at the given resolution. The connector element is shown in green. Enlarged views on the bottom row show the presence of only one connector element in the *Cr*NPC (red box) that is absent at the cytoplasmic ring of the *Cr*NPC (blue box) and the presence of two connector elements and duplicated Y-complexes in the *Hs*NPC (purple box). Abbreviations: cytoplasmic ring (CR), inner ring (IR), nuclear ring (NR).

We found that two key features of outer ring architecture are missing from the *Cr*NPC. First, the cytoplasmic ring contains only eight Y-complexes, half the number found in the *Hs*NPC. The Y-complex duplication is missing from the *Cr*NPC cytoplasmic ring. This explains why the cytoplasmic ring has less density than the nuclear ring, which contains 16 Y-complexes that are in rotational register with the 16 Y-complexes of the *Hs*NPC. *C. reinhardtii* thus has a total of only 24 Y-complexes, which are asymmetrically distributed across the nuclear envelope plane (16 in the nuclear ring, 8 in the cytoplasmic ring), in contrast to any previously proposed model of NPC scaffold architecture (Fig. 3, Video 1). This oligomeric state is consistent with the finding that metazoan-specific Nup358, which is required for linking the inner and outer Y-complexes of the cytoplasmic ring in humans ^27^, is absent in algae (Supplementary Fig. 6).

Second, the connector density attributed to *Hs*Nup155 in the *Hs*NPC^27^ is missing from the cytoplasmic but not the nuclear side the *Cr*NPC (Fig. 3b). This is surprising because this connector is the only rigid structural element that connects the inner ring to the outer rings in the *Hs*NPC. We therefore inspected the contact points between the inner and cytoplasmic rings of the *Cr*NPC. We found that contact is made by densities attributed to large scaffold Nups (Nup188 or Nup205) in the cytoplasmic ring of the *Hs*NPC (Fig. 3). We conclude that although the *C. reinhardtii* Y-complexes of the cytoplasmic ring arrange in a head-to-tail fashion similarly to humans, neither the oligomeric state nor the connection to the inner ring is conserved between the alga and humans.

### Subcomplexes of the cytoplasmic filaments differ between the algal and human NPCs

The Nup214 subcomplex (Nup159 subcomplex in fungi) is a key player in the remodeling and export of messenger ribonucleoprotein particles^1^. It is a major component of the cytoplasmic filaments that decorate the NPC scaffold at the cytoplasmic ring. In both the *Cr*NPC and *Hs*NPC, we observed characteristic densities extending from the cytoplasmic ring towards the central channel. However, the two densities are considerably different. The density protruding from the *C. reinhardtii* cytoplasmic ring is relatively large (black arrowheads, Fig. 1c) and would be consistent with previous analysis based on subtomogram averaging and cross-linking mass spectrometry that has associated the Nup159 subcomplex with the small arm of the Y-complex^4,29,30^. The corresponding density protruding from the human cytoplasmic ring is smaller (green arrowheads, Fig. 1c). This is may be due to flexibility or different subcomplex oligomeric state, emphasizing species-specific differences of this rather poorly conserved NPC module. At the given resolution, neither the algal nor the human NPC density map can accommodate the dimeric yeast Nup159 subcomplex^29,30^. Taken together, this analysis suggests that not only the Y-complexes but also more peripheral subcomplexes are subject to extensive variation across the tree of life.

## Discussion

The evolution of the NPC is deeply rooted with the origin of eukaryotes. The protocoatamer hypothesis suggests that NPCs and trafficking vesicles arose from a common ancestor by divergent evolution^31^. Understanding the evolution of the NPC is therefore pivotal for addressing the origin of eukaryotic compartmentalization. Although most Nups were postulated to be ancient proteins^23^, it remains unclear to what extent the organizational principles of the NPC are preserved in subsequent eukaryotic lineages. Here, by comparing NPCs of species from two distant eukaryotic kingdoms, we find that the oligomeric state of the NPC can be vary substantially.

Our findings are derived from the *Cr*NPC structure obtained by *in situ* cryo-ET. Analysis of the *C. reinhardtii* genome reveals that this alga has orthologs of all necessary protein constituents of the major Nup scaffolding subcomplexes known from other species, and our systematic fitting analysis of the *Cr*NPC supports this conclusion. While we cannot exclude that the compositional variability of the *Cr*NPC extends even further (e.g. through unidentified Nups specific to algae or Nup paralogs not yet included in the current genomic sequence), the assignment at the level of subcomplexes already reveals striking features.

In particular, the density map reveals that the *Cr*NPC contains a high degree of asymmetric density, with a total of only 24 Y-complexes, highlighting the importance of asymmetric linker Nups that are required to connect scaffold Nups^32^. Interestingly, a recent biochemical and morphological study of the Trypanosome NPC suggested that its NPC structure may be highly symmetric^12^. Although one might hypothesize that only 16 Y-complexes were present in the outer rings of ancient NPCs, with a similar stoichiometry as proposed for the yeast NPC^33^, it remains unclear if the oligomeric state of the *Cr*NPC arose due to a loss or a gain of function. Since a highly similar mode of nuclear Y-complex duplication is found in algae (*C. reinhardtii*) and vertebrates (humans), we consider it likely that vertebrates have duplicated their cytoplasmic Y-complexes using protein-protein interfaces that had already evolved for the nuclear ring, but using the metazoan-specific Nup358 as a dimerizer^27^. Such oligomeric duplication events are frequently observed during the evolution of protein complexes ^34^.

The inner ring of the *Cr*NPC map is dilated in comparison to the *Hs*NPC map. The *Cr*NPC inner ring has the same diameter as the outer rings, which are horizontally stacked upon it. The inner ring’s rotationally symmetric spokes are distantly spaced, thereby forming relatively large peripheral channels that have been proposed to accommodate inner nuclear membrane protein import^35^. In this conformation, the head-to-tail connection of the outer ring Y-complexes might be important for restricting the maximal dilation of the pores. Are these species-specific differences, or could they be related to the NPC’s functional state? Independent cryo-ET structural analysis suggests that such elaborate conformational changes might also occur in vertebrates^36^. Constricted inner ring conformations have been observed not only in isolated *X. laevis* and HeLa cell nuclear envelopes but also within intact u2os cells^8^, while more dilated conformations were observed within intact HeLa cells^7^. Taken together with our data from intact *C. reinhardtii* cells, these findings suggest that both constricted and dilated conformations have physiological relevance. We speculate that not only the FG-rich regions, but also the scaffold of the NPC may be much more dynamic than anticipated. Previous studies have reported the dilation of isolated *X. laevis* NPCs upon treatment with chemicals such as *trans*-cyclohexane-1,2-diol and steroids^37,38^. Using atomic force microscopy, these studies found that the NPC central channel diameter can expand up to 63 nm, the same diameter that we observed in *C. reinhardtii.* How such massive conformational changes are structurally induced and potentially regulated, awaits further analysis. The local FG-Nup concentration within the central channel might change during inner ring dilation. It remains to be determined whether inner ring dilation has any effect on nucleocytoplasmic transport activity, such as the rates and size limits of the transiting substrates, or whether it is relevant for inner nuclear membrane protein import.

Using *in situ* cryo-ET enabled by cryo-FIB milling, we were able to identify major structural variations within the NPC. Our study therefore underscores the importance of structural analysis within the native cellular environments of divergent species to understand the breadth of NPC architecture and ultimately gain insights into both NPC function and evolution.

## Materials and Methods

### Cryo-ET

Cells were prepared for data acquisition based on procedures described in *Schaffer*, *M. et al.*^39^. Briefly, cells were blotted onto EM grids, which were plunge-frozen into a liquid ethane/propane mixture using a Vitrobot mark IV (FEI) and then transferred onto a cryo stage in a Scios (FEI) or Quanta (FEI) FIB/SEM microscope. Cells were thinned with a gallium ion beam and transferred into a Titan Krios transmission electron microscope (FEI) equipped with a K2 Summit camera (Gatan) for tomogram acquisition, as described in *Albert, S. et al.*^17^.

### *Cr*NPC structure determination

Tomogram reconstruction and subtomogram averaging of the *Cr*NPC is described in an accompanying study^17^. Briefly, 78 NPCs were picked from twice-binned tomograms. Particles were manually aligned for correct orientation of the cytoplasmic and the nuclear rings of NPCs. Initial average of the whole NPC was calculated, using PyTom^40^, by imposing eight-fold symmetry. The eight asymmetric units of the indiviual aligned NPCs were extracted, yielding 624 asymmetric units. Alignment and averaging of these asymmetric units were carried out using the AV3/TOM packages as described^41^. After few iterations on the level of asymmetric units, masks specific to the cytoplasmic, nuclear and the inner ring of asymmetric unit were used to further align each of those parts respectively (as reported in^4^).

### Identification of *C. reinhardtii* Nups

The *C. reinhardtii* Nups were identified by retrieving predicted Nup sequences from the Phytozome platform ^42^ based on annotations or by BLAST ^43^ searches against the database of predicted *C. reinhardtii* proteins and the genomic sequence at Phytozome using human and plant Nups as queries. All identifications were confirmed using reverse BLAST searches (using the predicted Nups as queries) searches against a non-redundant protein database and by domain mapping using the HHpred server ^44^ to ensure that the identified genes are bona fide Nup orthologs rather than more remote homologs from other families (e.g. vesicle coat proteins).

### Assignment of subcomplexes within the *Cr*NPC map

To assign densities of the *Cr*NPC map to specific subcomplexes, a hierarchical fitting procedure (Supplementary Fig. 3) was applied as follows:

#### i) Unbiased global fitting

An unbiased global fitting approach was performed using structural models of various human subcomplexes derived from previously published structures^5,27^. The Y-complex was complemented with an structure (PDB ID: 4XMM)^45^. Because of the lower resolution of the *Cr*NPC map, all models were low-pass filtered to 30 Å. The resulting model maps were then independently fitted into the *Cr*NPC cryo-EM density using global fitting as implemented in UCSF Chimera^46^. All fitting runs were performed using 1,000,000 random initial placements, correlation about the mean as a fitting metric, and requiring at least 30% of the model map to be covered by the *Cr*NPC density envelope defined at low threshold. For each fitted model, this yielded 5,000-25,000 fits after clustering.

#### ii) Assignment of statistically significant fits

For each fitting run, the statistical significance of the fits was calculated as described previously^4,27^. All non-redundant statistically significant fits were placed in the model, leading to assignment of three copies of the inner ring subcomplexes. These three fits reproduced the arrangement observed in the inner ring of the *Hs*NPC, reinforcing the confidence in the fits.

#### ii) Assignment of the remaining densities by filtering top scoring nonoverlapping fits

To assign the remaining densities, for each fitting run of the inner ring protomers and outer ring Y-complexes, the top ten fits were selected and filtered according to following criteria: 1) overlap with the *Cr*NPC map was at least 60%, 2) the fits did not clash with the statistically significant fits already placed within the map, 3) the fits did not significantly overlap with the membrane density. This procedure led to a single solution of non-clashing fits including four copies of the protomer in the inner ring and three copies of the Y-complex in the outer rings (two in the nuclear ring and one in the cytoplasmic ring). All fits reproduced an overall arrangement that resembled the *Hs*NPC, increasing the confidence in the fits.

#### iv) Tentative assignment of Nup188/205 and the Nup155 connector

Analysis of the difference density between the resulting model and the *Cr*NPC map revealed characteristic unassigned densities on the nuclear side of the *Cr*NPC that matched the positions of Nup188/205 and the connector Nup155 in the *Hs*NPC. Based on both the shape and positional similarity, these densities were tentatively assigned as Nup188/205 and Nup155. Because the shape of Nup188 and Nup205 crystal structure is similar at 30 Å resolution, the densities could not be unambiguously assigned to one of the two Nups.

#### v) Optimization of the fits

Visual inspection of the fits indicated conformational differences between the fitted human subcomplexes and *Cr*NPC densities, especially in the stem region of the Y-complex. Therefore, the fits were optimized by local re-fitting of individual subunits or domains. It must be noted that due to the lower resolution of the *Cr*NPC map, the final fits should not be interpreted at atomic resolution; the flexible fitting merely aids in the assignment of densities and segmentations. Finally, Nup37 and the Elys β-propeller, which lacked corresponding densities in the *Cr*NPC map and were not identified in the *C. reinhardtii* genome, were removed from the Y-complex fits.

In addition to the above procedure, several validation runs were performed using the entire inner ring asymmetric unit (which led to a statistically significant hit, Supplementary Fig.4).

## Acknowledgments

We thank Drs. Felix Willmund and Jacob Musser for advice. Christian Zimmerli, Marc Wehmer, Drs. Elizabeth Villa, Eri Sakata and Matteo Allegretti are acknowledged for help with the preparation of this manuscript. S.M. and J.K. were supported by the EMBL Interdisciplinary Postdoc Programme under Marie Curie COFUND Actions. W.B. acknowledges funding from the Max Planck Society and Deutsche Forschungsgemeinschaft excellence clusters CIPSM and SFB 1035. M.B. acknowledges funding by EMBL and the European Research Council (309271-NPCAtlas).

## Author contributions

FIB milling: MS; cryo-ET: MS, BE, SA; structural analysis: SM, SA; structural modeling: SM, JK; bioinformatic analysis: JK; project management: JP, WB, BE, MB; paper writing: SM, JK, WB, BE, MB.

## Data availability

Cryo-EM maps of the *C. reinhardtii* cytoplasmic, inner and nuclear rings will be deposited into the EMDB. The cryo-EM map of the human inner ring has been previously published (EMD-8087).

